# GLP-1 agonism alters local field potential in the lateral septum and alters operant behavior in rats

**DOI:** 10.64898/2026.04.19.719508

**Authors:** Isabella R. Culshaw, Owen D. Jones, Ryan D. Ward, Robert G.K. Munn

## Abstract

GLP-1 agonists are an emerging treatment for disorders of consumption. They are most prominent as treatments for obesity, but recent literature suggests that they are effective at reducing the consumption of all types of hedonic substances. This clearly suggests a central, cognitive, mechanism rather than a peripheral mechanism or an interaction with a single signalling pathway, but the specific site or sites for this mechanism remain to be discovered. Candidate brain regions for this reward-modulating activity have a relative paucity of GLP-1 receptors, with the exception of lateral septum, which expresses an abundance of them. In these experiments we recorded local field potential from lateral septum while animals received either saline control or the GLP-1R agonist liraglutide. We find that liraglutide significantly reduced the power of both high-frequency oscillations and theta rhythm in the lateral septum, suggesting that GLP-1R agonism changes how lateral septum communicates with its network. In addition, we show that liraglutide causes animals to wait longer to respond for reward in a differential reinforcement of low rates paradigm. Together, these results suggest that a primary region in the control of the anticonsumptive action of GLP-1 agonists is the lateral septum, and that the coding of reward by this region is a central node in the network responsible for cognition about and behaviour with respect to reward.

## Introduction

Glucagon-like peptide 1 receptor (GLP-1R) agonists have quickly gained popularity for the treatment of diabetes and obesity. The GLP-1 system is well known to influence metabolic homeostasis of blood sugar, but it also has been shown to regulate feeding and therefore reduce bodyweight; this is thought to be mediated via its impact on food reward (Alhadeff et al., 2012; Dossat et al., 2013). It is becoming clear that agonism of this receptor system significantly influences circuits involved in motivation, reward-seeking and addiction-like behaviour, as GLP-1R agonism reduces the consumption of reinforcing substances generally (Egecioglu et al., 2013a; Shirazi et al., 2013; Harasta et al., 2015; Sørensen et al., 2016), ruling out interaction with a single pharmacological pathway, and suggesting a central mechanism (Srinivasan et al., 2025). It is therefore of interest to examine reward circuitry in the central nervous system as a substrate for the action of GLP-1 agonists. Activity in dopaminergic areas associated with reward such as the ventral tegmental area are modified by GLP-1R agonism (Merkel et al., 2025) but contain few GLP-1 receptors, implying control of the GLP-1 effect lies outside these traditional reward areas.

The Lateral Septum (LS) is critical for the perception of reward (Harasta et al., 2015); stimulation of the LS is highly rewarding (Olds and Milner, 1954; Sheehan et al., 2004) and LS neurons are highly responsive to drugs of abuse (Sheehan et al., 2004). The LS has also been implicated in addiction and relapse-like behaviours to drugs including methamphetamine, cocaine, and morphine (Le Merrer et al., 2007; Zahm et al., 2010; Cornish et al., 2012). In addition, it shares strong bidirectional connections to the Ventral Tegmental Area (VTA) (Risold and Swanson, 1997), linking it to systems which govern the expression of reward-directed behaviour. Unlike many regions involved in the reward circuit, the LS is rich with GLP-1Rs (Merchenthaler et al., 1999), particularly expressed by GABAergic neurons within LS (Luo et al., 2011). This expression is conserved across multiple species, including mice (Harasta et al., 2015; Jensen et al., 2018), rats (Terrill et al., 2016), and non-human primates (Heppner et al., 2015). GLP-1 signalling in LS has also been shown to attenuate reward-directed behaviour (Harasta et al., 2015).

LS is a central node in a network connecting a variety of regions across the brain responsible for forming a neural representation of the world. The most critical connection in this circuit is between the hippocampus and LS. Decades of research has demonstrated that neurons in hippocampus reflect a nuanced representation of egocentric space and time LS receives strong, almost monodirectional input from hippocampus (Rizzi-Wise and Wang, 2021). A population of GABAergic neurons within LS project to the VTA linking contextual information from the hippocampus with reward in the VTA. This circuit underlies context-dependent aspects of reward learning and reinstatement of cocaine-seeking: a model of relapse (Luo et al., 2011). Together, LS and hippocampus are part of the septohippocampal system. Connectivity between hippocampus and LS is temporally coupled through coincident fast events (Tingley and Buzsáki, 2020). These high-frequency sharp-wave ripples (SWRs, termed high-frequency oscillations (HFOs) in lateral septum) in the hippocampus are critical for learning and memory retrieval (Ego-Stengel and Wilson, 2010; Joo and Frank, 2018; Fernández-Ruiz et al., 2019; Pfeiffer, 2022). Hippocampal SWRs are temporally linked with LS HFOs (< 50 ms) (Tingley and Buzsáki, 2020) and this has led to the hypothesis that these events are causally linked; it is thought that SWRs drive downstream LS HFOs (Tingley and Buzsáki, 2020). This is likely a mechanism by which contextual information is coupled with reward-related information, and the combined output is then relayed to the VTA. LS is therefore optimally positioned to link GLP-1R activation with reward-related electrophysiological events and behaviour.

GLP-1R induced electrophysiological changes are hypothesised to result in a dampening of incentive salience, expressed behaviourally as reduced impulsivity and improved behavioural inhibition. A task which measures these facets of motivated behaviour is the differential reinforcement at low rates of responding (DRL) task. This is an operant paradigm which assesses behavioural inhibition, reward sensitivity, and the ability to regulate responding over time (Jarrard and Becker, 1977; Gage et al., 1979; Doughty and Richards, 2002; Munn and McNaughton, 2008; Eckard and Kyonka, 2018). Current evidence indicates that GLP-1R agonists reduce wanting (reduced drug or food seeking behaviour) but do not obviously affect liking (hedonic reaction), the key component of the DRL task is the use of a forced time delay which dissociates ‘wanting’ from the ‘liking’ aspect of motivated behaviour by requiring a subject to inhibit a previously shaped operant response (lever pressing) to obtain a reward. If a subject ‘wants’ the reward more, the average time between responses are likely to be reduced as they are less able to inhibit the behaviour, decreasing performance and vice versa, It has previously been established that there is a strong link between ‘wanting’ and impulsivity, which can then increase problematic substance use behaviour (File et al., 2022).

Despite emerging evidence that LS is likely a crucial substrate for GLP-1, the effects of systemic GLP-1R agonists on LS function are unknown. It is also unknown which specific aspects of reward-related behaviour these drugs affect. This study therefore aimed to examine the effects of the GLP-1R agonist liraglutide on the electrophysiology of LS, and to investigate liraglutide’s effects on behavioural inhibition via the DRL task.

## 2. Methods

### Subjects

All procedures were approved by the University of Otago Animal Ethics Committee and conducted in accordance with institutional guidelines. Male (electrophysiological experiments n = 3) and female (behavioural experiments n = 12) Sprague-Dawley rats were sourced from Otago University’s Breeding Research Facility when aged 8-11 weeks old. All animals were given access to food and water *ad libitum* prior to any manipulations and kept in a colony room with a 12 hr light-dark cycle (lights on 7 am), maintained at 22°C□. Rats habituated to laboratory housing for at least 2 weeks after arrival before any experimental manipulations. During this time animals were habituated to daily handling.

### Drug treatment

Liraglutide 6 mg/ml injection pens (Saxenda®, Novo Nordisk Inc.) were obtained (3 ml solution per pen, pH ∼8.15; NovoNordisk) Pens delivered measured doses of 0.6, 1.2, 1.8, 2.4and 3.0 mg. A dilution was therefore performed to produce injection volumes in the 0.2-1ml range. A 0.6 mg dose (0.1 ml) was diluted 1:100 in physiological saline, giving a final concentration 0.06 mg/ml liraglutide). Pens were kept refrigerated (2-8 °C) when not in use. Syringes were allowed to come to room temperature before injecting rats, to prevent discomfort or shock. Animals were administered 0.03 mg/kg or 0.06 mg/kg depending on experimental group.

### Electrophysiology

#### Subjects

Experiments used *n* = 3 male rats aged 3-6 months (400-500 g). Rats were group housed in an individually vented cage (49 x 31 x 26.5 cm) until surgery, after which they were housed individually. Prior to surgery and testing, all rats were placed on a food deprivation regimen to maintain body weight at ∼85% of free-feeding weight (∼15-20 g chow per day). Experimental sessions (*n* = 6; *n* = 1 from one rat, *n* = 2 from another, and *n* = 3 from the third) were conducted between 11 am - 3 pm.

#### Habituation

Prior to habituation and recording, Kellog’s Coco Pops were distributed in home cages to familiarise animals with this reward and facilitate exploration behaviour in the open field arena. Animals were initially habituated to the open field apparatus (A square black plastic tub, measuring 100 x 100 x 50 cm). over a week with daily sessions lasting < 1 hr with lights off in the experimental room. Rats were habituated until they exhibited consistent exploratory behaviour when placed into the field and did not exhibit anxiety or fear behaviours (e.g. hunching, freezing).

#### Microdrives and stereotaxic surgery

Prior to experiments, subjects underwent stereotaxic surgery for implantation of electrodes in LS (Munn et al., 2022). Implants were custom built 16-channel microdrives, made up of four tetrodes. Implants were directed toward the caudo-dorsal portion of LS at coordinates: Bregma +0.05, midline −0.5 mm (Wirtshafter and Wilson, 2020)and 4 mm deep. This placed electrodes immediately superior to the LS. All animals received post-operative care, with heat and fluid support (5 ml physiological saline administered subcutaneously (s.c.) on each side of the abdomen), as well as additional pain-relief (Carprofen, once daily, 3 days). A post-surgery recovery period of 2-weeks was observed before conducting any recordings.

#### Electrophysiological Recording Apparatus

Data were acquired using the DacqUSB (Axona Inc.) system. LFP data were digitally amplified by a preamp and sampled simultaneously at low (250 Hz) and high (4.8 kHz) frequency optimised to capture theta and HFO activity, respectively. Animal positions in the X-Y plane were tracked using an overhead camera connected via coax to the DacqUSB system and sampled at 50 Hz. LEDs on the head mounted cable also assisted with tracking animal position.

#### Electrophysiological recordings

Daily recordings screened for the presence of dominant theta-band oscillations in the LFP indicating the electrodes were in LS. If theta was absent, electrodes were advanced ∼25 μm and animals were returned to their home cage until the next day. If theta frequency was present, animals moved on to an experimental recording session. Animals were weighed at the beginning of experimental days to calculate drug dosage. Subjects were injected s.c. with saline vehicle for a control session, then placed back into the home cage for a 60 min waiting period to match the drug session. After this, rats were moved into the open field and allowed to freely explore for 20-40 min during electrophysiological recording. Following this they were injected with 0.06 mg/kg liraglutide and placed back into the home cage for 60 min. Liraglutide exhibits rapid kinetics, with behavioural effects seen after as little as 10 mins (Herman and Schmidt, 2024). In addition to its rapid action, it has a half-life of 13 hours in humans (Drucker et al., 2010) and approximately 4 hours in rats (Sturis et al., 2003). which ensures that the 60 min wait period for drug effect would not lead to a premature wash-out. Following this wait period, recording began again. From the x-y position of the LEDs, movement speed was calculated and the time spent immobile (speed < 1 cm/s) was calculated. Between sessions, the open field was wiped down with 80% EtOH.

#### Data processing and analysis

LFP data were analysed using a combination of existing and custom-written functions in MATLAB (v.2025a, Mathworks, Inc), and the chronux toolbox (Mitra, 2007) (chronux.org). Spectral power of recorded LFP was calculated using the chronux function `mtspecgramc` with a moving window of 2 seconds and a step of 0.25 seconds. To determine theta frequency, LFP was bandpass filtered between 5-12 Hz using the bandpass function and then subjected to a Fourier transform using the function `fft.` The `hilbert` function was used to determine the instantaneous phase and frequency. Average frequency per session was calculated from the instantaneous frequency of each sample.

##### Detection of high-frequency oscillations

The method we used to identify high-frequency oscillations was derived from previous studies identifying sharp-wave ripples in the hippocampus (Karlsson and Frank, 2009; Jadhav et al., 2012; Yu et al., 2017). Briefly, LS LFP was filtered in the ripple band between 150-250Hz. The envelope of the resulting filtered LFP was obtained by Hilbert transform and smoothed with a gaussian kernel (s = 2ms). Segments of the resultant filtered and smoothed LFP were considered a HFO if the amplitude was > 3 SD above the mean amplitude of the filtered signal for a duration of at least 20 but not more than 250ms. Ripple events were isolated at the power of these events was calculated using a continuous wavelet transform with MATLAB function `cwt` with a wavelet built with `cwtfilterbank` with frequency limits 80-200Hz. The cumulative probability power curves were compared via two-sample Kolmorgov-Smirnov test. Theta frequency and running speed data were analysed using paired t-tests (alpha = 0.05). All data are presented as mean ± SEM.

### Behavioural testing

#### Subjects

Twelve female Sprague-Dawley rats (8-11 weeks upon arrival at the laboratory, 190-250 g at the beginning of the experiment) were used. To increase motivation for food rewards, rats were placed on food deprivation and maintained at 90% of their free-feeding weight for experiments. Rats were housed in pairs to avoid social isolation stress. Rats received regular chow pellets every day at the same time during food deprivation (∼ 1:00 pm for first session, 2:30 pm for second session; in addition to food consumed during experiments).

#### Behavioural apparatus

Six identical Med-Associates operant chambers were programmed to run FR-1 (lever-press association), DRL-5, -10, and -15 tasks. Each rat was assigned an operant chamber for the entirety of habituation, training, and experimental sessions. Chambers were wiped down with 80% EtOH before and after each session.

#### Pretraining

Prior to training, rats were given rewarding pellets in home cages (10-20 g daily, 2 days) to facilitate consumption during experiments. Rats were randomly split into 2 experimental groups of 6 on a by-cage basis to minimise separation from cagemates. Animals were habituated to operant chambers with sessions of 20 min with the fan running (no lever or house light) every day for 5 days. We used a validated auto-shaping method (Deane et al., 2017; Meighan et al., 2021) to train rats to nose-poke for reward. Next, subjects were trained to associate lever-pressing with pellet delivery. The left lever was extended into the chamber and, if the rat pressed the lever within 10□s, a pellet was delivered into the receptacle. If the lever was not pressed within 10□s, the lever was retracted and the next trial began. The criterion for progressing was met when rats were lever-pressing on more than 80% of the 60 trials.

#### Training

Animals began training on the DRL-5 task (5 sec must elapse between responses). For the first week of training, animals remained in the operant chamber until 60 rewards had been obtained (∼2 h per session). After this, sessions lasted for a maximum of 1 h. After ∼2 weeks, animals were reliably performing the DRL-5 at criterion (retrieving > 75% (45/60) of possible rewards), and they progressed to DRL-10 training for ∼7 days, until all animals were at criterion. Animals were then moved to the DRL-15 with session duration, training duration, and criterion as above.

#### DRL-15 with Acute Liraglutide

We investigated the impact of acute liraglutide administration on performance in the DRL-15 task. All rats received SC injections of physiological saline 60 min prior to completing a single DRL-15 session (Session A). After 1 week, rats were injected with 0.03 mg/kg liraglutide and, after a 60 min waiting period, completed another single DRL-15 session (Session B). One week later, injections of 0.06 mg/kg liraglutide (Session C) were administered 60 min before the DRL-15 sessions. Between experimental sessions, all subjects underwent 1 DRL-15 session per day with no drug to maintain performance at criterion. Injections were administered at the same time every day (∼11:00am for the first session and 12:30pm for the second session). R rats were returned to their home cages for the 60 min post-drug administration period, prior to being moved to the experimental room and placed in operant chambers. Rats underwent DRL-15 sessions in batches of 6. Each animal therefore had 1 experimental session per day. This occurred at the same time every day during the light cycle (12:00 pm for first batch, 1:30 pm for second batch). After the first DRL-15 experiment, there was a 7-d drug washout period during which all animals underwent a daily DRL-15 session to maintain behavioural readiness.

#### DRL-15 chronic

To investigate the impact of chronic liraglutide administration, all subjects received daily injections of 0 mg/kg (saline), 0.03 mg/kg, or 0.06 mg/kg liraglutide across a period of 15 d days 1-5 0.9% saline, days 6-10 0.03 mg/kg liraglutide, days 11-15 0.06 mg/kg liraglutide). Each day, all animals completed one DRL-15 session. After completing each session rats were returned to their home cages and received their regular daily chow meal.

#### DRL-15 behavioural measures

For the inter-response time per opportunity for reward (IRT/OP) measure, responses were binned according to specific time intervals. The IRT/OP for each bin was then defined as the probability of a lever press occurring within that bin given that a subject had waited at least as long as the start time of the bin in question after the last response (Anger, 1956; Tonkiss and Rawlins, 1992). For example, if responses were binned into 5 s intervals, the IRT/OP for the bin 5-10 s was the number of responses made between 5 and 10 s after the last response, divided by the sum of the number of responses in bin 5-10 and all of the future remaining time bins (i.e. 10-15, 15-20 etc.). The IRT/OP for a given bin (x) was therefore represented as:

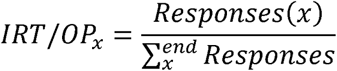

#### Statistical analyses

All data collected during behavioural testing were analysed using a combination of MATLAB 2025a (Mathworks Inc.), Prism v10.1.1 (GraphPad Software LLC), and JASP v 0.19. In acute experiments, all measures apart from IRT/OP were analysed using one-way ANOVA, with dose as a repeated measure. IRT/OP was analysed using a repeated measures ANOVA with polynomial contrasts for the within-subjects variables bin and drug. IRT/OP was analysed using a repeated-measures ANOVA with polynomial contrasts for dose and bin (within-subjects) and with day of testing extracted as between-subjects factors. For chronic DRL-15 number of responses data were analysed using a repeated measures ANOVA with the between-subjects factors day and drug.

### 3. Results

### 3.1 0.06mg/kg liraglutide reduces the power of high frequency oscillations in the lateral septum

High frequency oscillations (HFOs; Figure 1D) were recorded from the LS of n = 3 rats across n = 6 recording sessions (n = 3 from one rat, n = 2 from another and n =1 from the third). When animals were administered 0.9% saline, 575 ripple events were detected. When animals were given 0.06mg/kg liraglutide, 1036 events were detected. HFO events were more likely to have lower power when animals were given 0.06mg/kg Liraglutide compared to control (Figure 1A-C two-sample Kolmogorov-Smirnov test, D = 0.114, p = 1.145 x 10^−4^).

**Figure 1.**
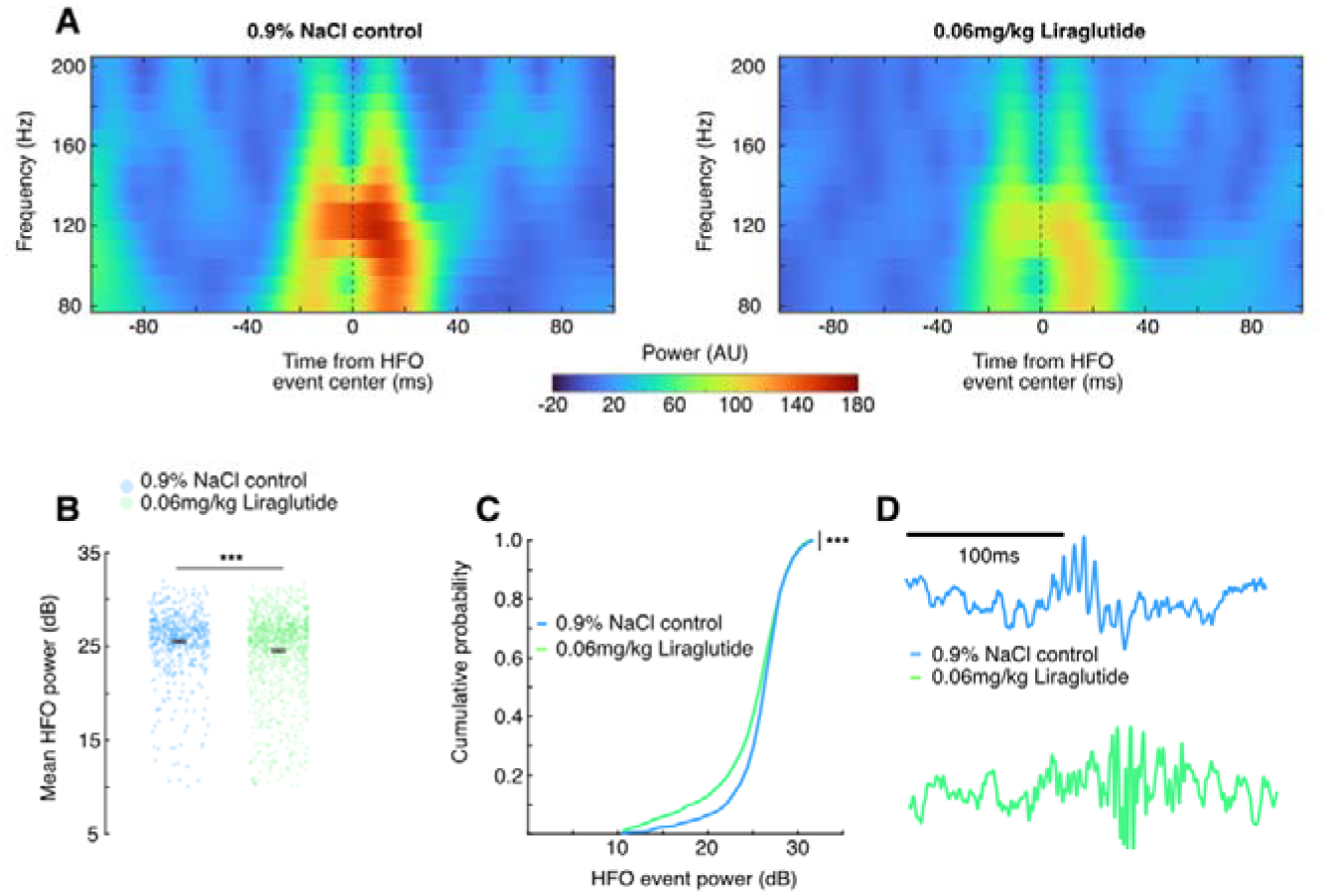
Liraglutide reduces the power of HFOs in the lateral septum. **A,** the mean power over all HFO events recorded while animals were administered either saline control (left) or 0.06mg/kg Liraglutide (right). The 200ms time window for each is centered in the middle of each event, and this is illustrated with a black dotted line. **B,** Decibel-transformed power of HFO events recorded when animals were administered saline control (blue) or 0.06mg/kg Liraglutide (green). The mean and ± SEM are shown as a red point with black bars. **C,** the cumulative probability that events in LS are a given power when animals are administered saline control (blue) or 0.06mg/kg Liraglutide (green). **D,** Examples of HFO events in LS in the control (blue) and 0.06mg/kg Liraglutide (green) conditions. ***p < 0.01

### 3.2 0.06mg/kg Liraglutide reduces the power, and increases the frequency of theta rhythm in the lateral septum

We next examined spectral power in common frequency bands associated with activity in the septohippocampal formation. 0.06mg/kg liraglutide appeared to cause an increase in low-frequency delta-band (0.5-4Hz) power, but this difference was not significant (Figure 2-C, mean power (dB) ± SEM, control = 65.0 ± 0.58, liraglutide = 63.9 ± 2.18, t(5) = 0.41, p = 0.70, n.s.). Although 0.06mg/kg liraglutide appeared to reduce the power of beta (13-30Hz) oscillations compared to control, there was no significant difference in power between these conditions (mean power (dB) ± SEM, control = 55.4 ± 0.51, 0.06mg/kg Liraglutide = 54.3 ± 0.75, t(5) = 2.05, p = 0.096, n.s.) The theta peak observed in the spectrum of rats given 0.9% saline was notably reduced when animals were given 0.06mg/kg Liraglutide, however (Figure 2C,D mean power (dB) ± SEM, control = 63.5 ± 0.72, liraglutide = 61.4 ± 0.77, t(5) = 2.82, p = 0.037). In contrast to the reduction in theta power, liraglutide increased the frequency of LS theta compared to control (Figure 2E, mean frequency (Hz) ± SEM, control = 9.28 ± 0.04, Liraglutide = 9.50 ± 0.05, t(5) = 3.02, p = 0.029). Theta amplitude and frequency are positively correlated with movement speed (or acceleration) (Kropff et al., 2021; Kennedy et al., 2022), so one explanation might be that Liraglutide makes animals move more quickly. Instead, we find that 0.06mg/kg Liraglutide significantly reduced the average movement speed of animals compared to control (Figure 2F, mean speed (cm/s) ± SEM, control = 15.46 ± 2.67, Liraglutide = 8.60 ± 1.60, t(5) = 5.29, p = 0.0032), making this explanation unlikely.

**Figure 2.**
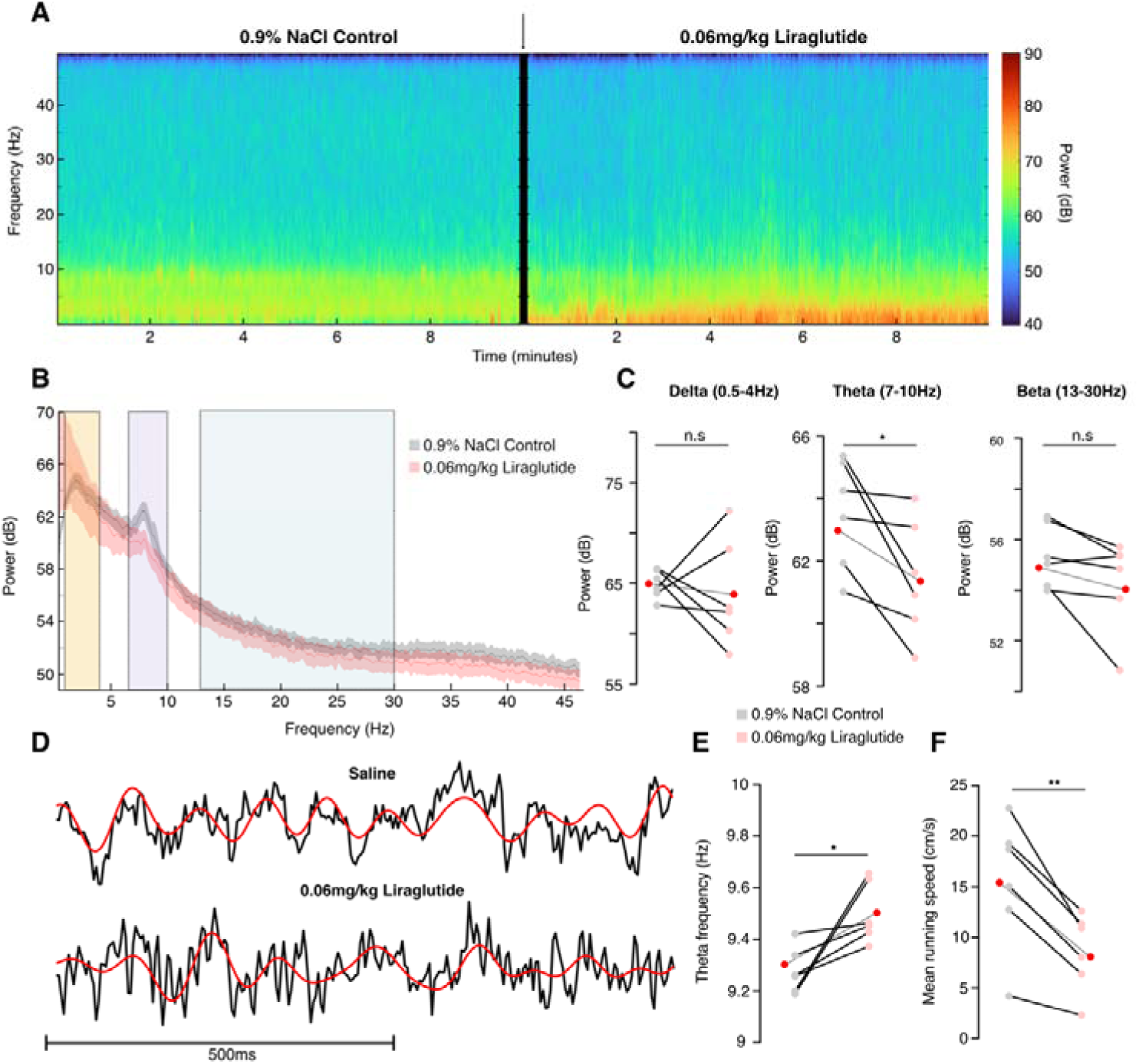
Liraglutide lowers the power and increases the frequency of theta rhythm in LS, while reducing speed. **A,** Mean decibel-transformed spectral power over all recording sessions when animals were administered saline control (left panel) and 0.06mg/kg Liraglutide (right panel). Time of injection is denoted with an arrow between the panels. **B,** mean (solid line) ± SEM (shaded region) decibel-transformed power from 0-50Hz over all recording sessions when animals were administered saline (grey) and 0.06mg/kg Liraglutide (red). Colored rectangles illustrate the frequency envelopes of delta (0.5–4Hz, orange), theta (7-10Hz, purple), and beta (13-30Hz, green) bands. **C,** Mean spectral power in the delta (left), theta (middle), and beta (right) bands. Mean power in each recording session is illustrated by grey points, and each saline recording session is joined with the corresponding Liraglutide session by a solid line. The mean power over sessions is illustrated by red points joined by a dashed line. **D,** Example raw 1-second long LFP segments (black) with the theta bandpass filtered LFP overlaid (red). **E,** as in (C), but for theta frequency. **F,** as in (E), but showing the mean running speed in cm/s. *p < 0.05, **p < 0.01.

### 3.3 Acute and Chronic administration of Liraglutide improves performance on the DRL-15 operant behavioral task

When animals (n = 12) were acutely exposed to saline, 0.03 mg/kg liraglutide, or 0.06 mg/kg liraglutide we observed an initially high probability of response (Figure 3A), which decreased rapidly over the first 5 s, then progressively increased up until the 15 s time bin. This was followed by a steady decrease until the 30 s time bin. Overall, there was a significant linear interaction (drug x bin, lin x lin t(11,616) = 5.432, p<0.001). Figure 3B illustrates the mean IRT/OP (%) of subjects during each drug condition across 2 s time bins of the acute DRL-15 experiment. Each subject’s data for each bin were averaged across all sessions and days. Relatively few responses were likely to occur given that rats had waited for the earlier bin times (bins 0-10 s), the probability of a response occurring increased around the time of reward (between bins 11-18 s, given that an animal had waited this long), then from bins 20-24 s, a relatively lower probability of a response occurring here. Saline and drug groups had a low IRT/OP in the earlier bins, however, as time to reward approached there was an increase in IRT/OP in drug sessions compared to saline. Overall, there was a significant interaction (drug x bin, lin x quad, t(9,234) = 3.6, p = 0.006). Animals were more likely to wait longer in later bins with a higher IRT/OP in drug groups compared to saline.

**Figure 3.**
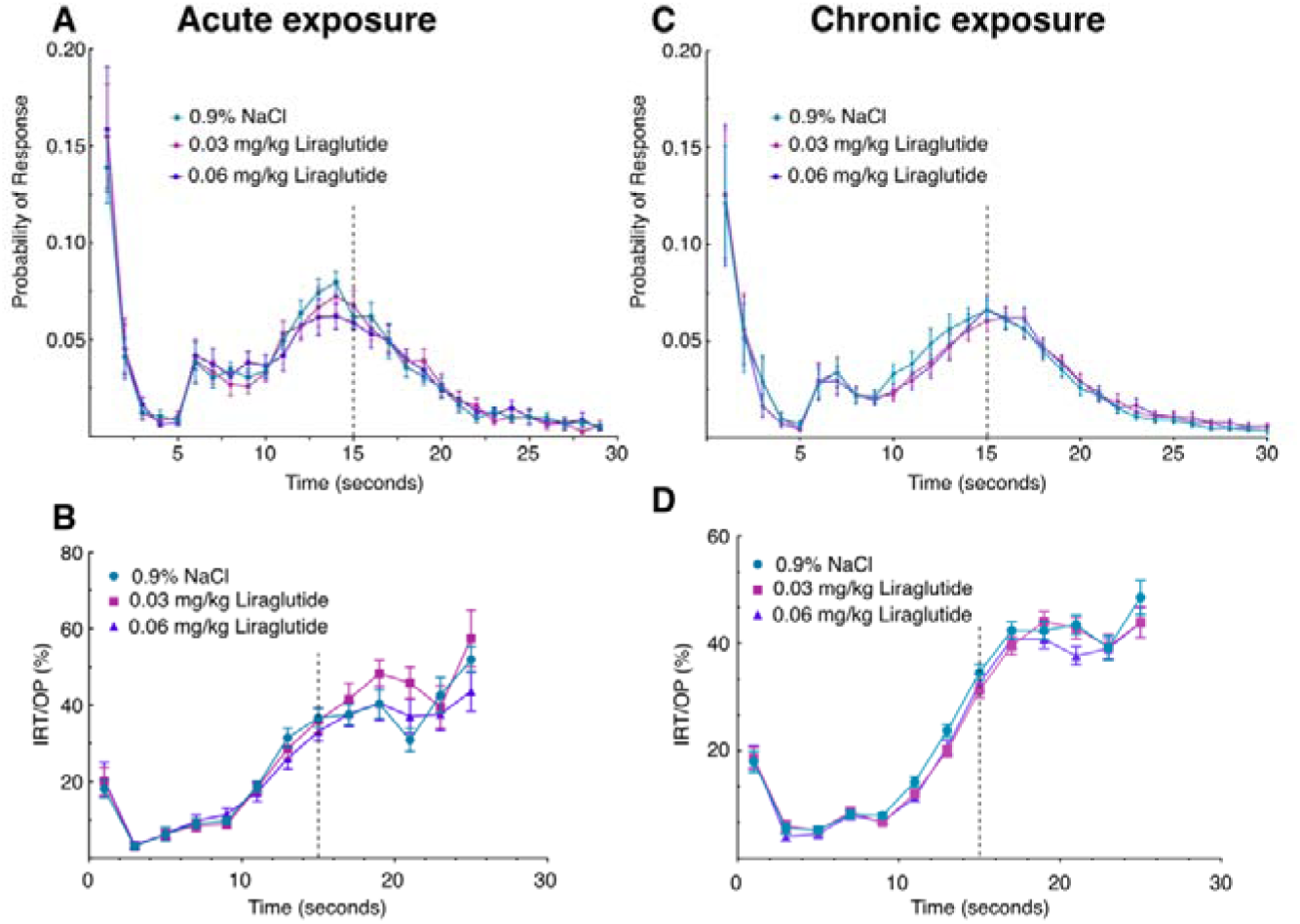
Liraglutide changes behavior in the DRL-15 task. **A,** the probability ± SEM of a response being made at a specific time (1-second bins) from last response when animals had acute exposure to saline (cyan), 0.03 mg/kg liraglutide (magenta), or 0.06 mg/kg liraglutide (purple). The time after which responses resulted in reward is shown with a blacked vertical dashed line. **B,** The IRT/OP for the data shown in (A). Data are binned in 2-second-long bins. **C,** as in (A), but averaged over sessions when animals were chronically administered drug or control. **D,** as in (B), but showing IRT/OP data from (C).

Next, when animals (n = 12) were chronically exposed to saline, 0.03 mg/kg liraglutide, or 0.06 mg/kg liraglutide, subjects exhibited a high probability of response (Figures 3C,4A) which decreased within the first 5 seconds. Probability was observed to increase up until the 15 s time bin (peri-reward period). Notably, when exposed to saline animals were more likely to respond prior to the reward time than when they were exposed to 0.03 mg/kg or 0.06 mg/kg liraglutide. This was followed by a decrease until the 30 s time bin. Overall, there was a significant interaction (drug x bin, lin x lin) t(11,638) = −3.575, p = 0.004). We next assessed the IRT/OP (%) of subjects during each drug condition across 2 s time bins of the chronic DRL-15 experiment (Figures 3D,4B). Data from each subject for each bin were averaged across all sessions and days. Given that an animal had waited for 2-10 s, there was a relatively low likelihood of responding for all groups. As the peri-reward period approached IRT/OP increased. After 20 s had elapsed, IRT/OPs began to decrease again across all drug conditions. Overall, there was a significant interaction effect (drug x bin, lin x quad, t(9,234) = 3.6, p = 0.006). Animals were more likely to wait longer in later bins with a higher IRT/OP in drug groups compared to saline. (Drug x bin, lin x lin, t(11,638) = −3.575, p = 0.004).

## Discussion

In this study, we sought to examine the impact of systemic GLP-1R agonist administration on LFP activity within LS, and to investigate its impacts on the ability to regulate operant responding, via the DRL task. For the first time, we have shown an effect of systemic liraglutide administration on LFP activity in LS. Acute liraglutide administration significantly reduced the mean power of HFOs within LS. Concurrently, we observed a significant increase in mean theta frequency accompanied by a significant reduction in mean running speed within the open field. In addition, we have presented evidence that systemic liraglutide enhances subjects’ behavioural inhibitory control during a DRL-15 schedule. Taken together, these findings show that GLP-1R agonism modulates LS activity and strongly impacts reward-related behaviour. While we cannot rule out roles for other brain structures, our results contribute to the growing theory that LS is a critical effector region in reward processing and is likely acting as a facilitator of subsequent reward-driven behaviour. High frequency oscillations are already thought to be a critical mechansim through which infomation is exchanged from a region with its communicating network. Given that LS is densely interconnected with the “traditional” dopaminergic reward circuitry (Rizzi-Wise and Wang, 2021; Besnard and Leroy, 2022; Tong et al., 2023) we propose that the modulation of LS HFOs by GLP-1R agonism is a core mechanism by which these drugs exert their effects on reward-related behaviour and cognition. Contributing to changes in reward-seeking and impulsive behaviour. It has already been established that LS is strongly implicated in drug-seeking and addiction-like behaviours (Le Merrer et al., 2007; Zahm et al., 2010; Cornish et al., 2012) This, in addition to the rich expression of GLP-1Rs here (Merchenthaler et al., 1999) by GABAergic neurons, and the ability of these same neurons to suppress reward-driven behaviours (Harasta et al., 2015) strongly supports our model, and the present findings are in reciprocal support. Whether changes in LS activity are causally linked to changes in behaviour needs to be investigated more directly, and this will be an important avenue for further study.

Typically, a positive correlation exists between theta frequency and running speed. In the present experiment, the opposite relationship was observed when subjects were administered liraglutide; animals ran more slowly, while theta frequency increased. The reduction in movement speed we observed is consistent with previous studies showing that systemic GLP-1R agonism can dose-dependently decrease locomotion (Bernosky-Smith et al., 2016). The alterations to theta frequency that we saw were therefore decoupled from locomotion. Instead, this likely reflects an effect on circuitry that subserves other aspects of behaviour.

Evidence suggests that excessive use of addictive drugs (e.g. alcohol, methamphetamine, nicotine etc.) can increase impulsive behaviour for reward (Jentsch and Taylor, 2001). Preclinical studies in animals and some human studies illustrate the opposite relationship: high trait impulsivity can be predictive of future drug-related behaviour (Anker et al., 2009; Winstanley et al., 2010). For example, in rodents higher levels of alcohol consumption are associated with higher levels of impulsive choice in a delay discounting task (Poulos et al., 1995). It is likely that this is a bidirectional relationship. In models of addiction, the dopamine response to drug-paired cues becomes potentiated. These cues then trigger greater drug craving and seeking (Winstanley et al., 2010). Previous work has shown that GLP-1R agonists suppress dopamine responses in the mesolimbic system to drugs such as nicotine, amphetamine, and cocaine (Egecioglu et al., 2013b; Sørensen et al., 2016). This may partly explain why GLP-1R agonists have shown promise in treating SUDs. Impairments in inhibitory control are implicated in many SUDs. Overall, these results suggest that the effects of GLP-1R agonists on reward behaviour likely stem from changes in incentive salience (motivation), which is expressed as the inhibition of impulsive responding seen in the DRL-15 task. Despite the use of a smaller cohort in the electrophysiology experiments, the impacts of this sample size were mitigated by our sampling method. The dosage levels used across both DRL-15 experiments, as well as the progressively increasing dose regimen used during the chronic DRL-15 experiment provided another methodological strength to this project. This regimen reflects the clinical use of prescribed GLP-1R agonists, with dosage typically escalating over time towards a higher, plateaued maintenance dose (Ratner et al., 2010). The lower dose (0.03 mg/kg) used is also comparable to that seen in human patients. Those involved in clinical trials for liraglutide are often > 27 BMI or within the 30-40 kg·m⁻² BMI range (Astrup et al., 2012; Davies et al., 2015), this corresponds to approximately 80–125 kg, depending on height of participants. The dosages for these studies were 1.2, 1.8, 2.4 and 3.0mg, which equates to a dose of approximately 0.03mg/kg.

## Conclusion

Liraglutide significantly reduced the mean power of HFO events and increased mean theta frequency recorded from LS, while simultaneously reducing mean running speed in the open field environment. In the DRL-15 task, acute liraglutide increased IRT/Ops as rewards approached in time, compared to saline. In addition, chronic liraglutide significantly reduced IRT/Ops. These behavioural effects may be mediated by changes in HFO and theta frequency activity within LS, however, further study is required to clarify this relationship. Our novel findings have strong translational applications and can inform future work investigating the use of GLP-1Rs in the treatment of compulsive disorders.

## Notes

### Competing Interest Statement

The authors have declared no competing interest.

